# Predictive coding networks capture human neural representations missing in supervised DNNs

**DOI:** 10.64898/2026.09.18.752626

**Authors:** Dirk Gütlin, Denise Kittelmann, Ryszard Auksztulewicz

## Abstract

Neuroscientific learning theories propose that the brain acquires knowledge by constructing internal world models. Supervised learning, the dominant approach in deep neural networks (DNN), relies on external category labels, making it difficult to reconcile with biological learning. There is an increasing trend towards more biologically valid approaches, such as predictive (minimize future surprise) or contrastive (minimize response to expected, maximize to unexpected inputs) objectives, but these approaches typically rely on pretrained DNN models that vary widely in architecture, size, and hyperparameters, making direct comparisons difficult. Here, we isolate the effect of learning by comparing small, identical networks trained with predictive, contrastive, and supervised learning objectives, as well as local (layer-restricted) vs. global (full backpropagation) learning. Using representational similarity analysis of a previously acquired EEG dataset, we show that brain representations after statistical learning are better modeled by a predictive local learning than supervised or contrastive learning, and that during learning, the brain attenuates category-specific representations while retaining predictive ones. We additionally show that predictive objectives explain brain variance that standard supervised DNNs do not, and that this variance is tied to predictive rather than stimulus or mismatch processing. These findings demonstrate a controlled approach for testing the algorithmic basis of learning and identify prediction as a core learning mechanism.

## Introduction

Forming internal representations of the relational structure of the environment is crucial for adaptive behavior. How such representations are acquired fundamentally relies on the brain’s learning mechanisms. In recent years, comparisons between biological systems such as the brain and artificial systems such as deep neural network (DNN) models have emerged as a powerful method to systematically address different questions about theories of brain computation including learning (Doerig et al., 2023; ***Kanwisher et al., 2023***; ***Schmitt et al., 2026***; ***Cichy and Kaiser, 2019***). Specifically, by manipulating network architecture (e.g., feedforward vs recurrent architecture), learning objective (e.g., supervised vs unsupervised) or learning mechanism (e.g., backpropagation), DNNs have provided insights into how these factors contribute to the emergence of neural representations in the brain. For example, by comparing different network architectures, previous work revealed the role of recurrent processing in shaping representational dynamics across the ventral visual stream during object processing (***Kietzmann et al., 2019***). Similarly, recent work comparing different learning objectives has demonstrated that unsupervised and self-supervised networks exhibit representations that align more closely with those observed across the ventral visual stream (e.g., ***Kataoka et al., 2025***; ***Prince et al., 2024***; ***Konkle and Alvarez, 2022***). Despite these advances, most successful DNN models rely on backpropagation, a learning algorithm that has been argued to only loosely reflect biological learning mechanisms in the brain (***Whittington and Bogacz, 2019***; ***Lillicrap et al., 2020***; ***Stork, 1989***). Specifically, backpropagation updates model parameters by recursively calculating gradients of the output error for each layer and propagating them backwards through the network hierarchy (Rumelhart et al., 1986a). Furthermore, many DNNs are optimized using supervised learning, relying on externally provided labels to guide learning (***LeCun et al., 2015***). However, unlike supervised learning, biological learning is generally thought to occur without access to explicit ground-truth labels or externally provided teaching signals. Instead, learning may largely emerge from sensory experience itself (***Sherman et al., 2020***). Building on these limitations, research has increasingly focused on developing biologically inspired learning algorithms in DNNs informed by biological mechanisms and neuroscientific theories of learning (e.g., ***Whittington and Bogacz, 2017***; ***Lotter et al., 2016***; ***Millidge et al., 2024***; ***Hinton, 2022***; ***Tscshantz et al., 2023***; ***Schmidgall et al., 2023***).

Frameworks such as Predictive Coding (PC) propose that the brain learns by iteratively generating predictions and updating internal models of the world through mismatch signals, i.e. signals elicited by mismatches between predicted and received input (***Friston, 2005***). Within this framework, both predictive and mismatch signals are thought to play a central role in learning. Consistent with this view, a large body of empirical evidence has revealed insights into the dynamic roles of predictive and mismatch signals over the course of learning (e.g., ***Aitken and Kok, 2022***; ***De Lange and Press, 2026***; ***McDermott et al., 2026***). More broadly, accumulating evidence has shown that learning modifies neural representations in the brain over time (***Greco et al., 2024***; ***Chiossi et al., 2025***; ***Poort et al., 2015***; ***Sherman et al., 2022***). However, how different learning principles shape the formation and modification of these neural representations remains poorly understood.

PC-inspired models provide a computational framework for studying these questions by implementing learning principles derived from the PC theory (***Whittington and Bogacz, 2017***; ***Millidge et al., 2024***, 2021). Rather than relying exclusively on globally propagated error signals, these models update their internal representations through locally computed errors, thereby providing a biologically motivated alternative to backpropagation (***Whittington and Bogacz, 2017***; ***Enan et al., 2025***). While various PC-inspired learning algorithms and specific predictive coding networks (PCNs) have been proposed that differ in their computational formulation, they are unified by a common predictive objective that drives learning through the minimization of prediction errors. Previous work has shown that PC-inspired learning algorithms achieve performance comparable to supervised DNNs on smaller tasks, although they are less commonly adopted in machine learning and more difficult to scale than conventional DNNs (***Whittington and Bogacz, 2017***; ***Millidge et al., 2021***). PC-inspired algorithms can also improve aspects such as catastrophic forgetting (***Lee et al., 2022***) or brain similarity (***Guo et al., 2025***). Similarly, a growing body of evidence suggests that representations learned by PC-inspired DNNs neural representations align more closely with neural representations in the brain than those learned by supervised DNNs (***Guo et al., 2025***) and exhibit less degradation in representational alignment across the cortical hierarchy (***Leutenegger, 2026***).

While these findings provide promising evidence consistent with theoretical proposals that PCNs might capture biological learning dynamics better than supervised trained DNNs, these comparisons often involve models that differ in architecture, scale, or training regime. Consequently, it remains difficult to attribute differences in model–brain correspondence specifically to the learning mechanism. However, pinpointing the specific contribution of learning mechanisms in these models is particularly important if DNNs are to provide mechanistic insights into biological learning. Recent work addressed this issue using small, architecturally identical RNNs to simulate the effects of different learning objectives and mechanisms and systematically tested to what extent predictive and contrastive PC-inspired learning objectives, compared with supervised objectives, exhibited key PC-like learning dynamics, such as prior expectations, mismatch responses and semantic representations (***Gütlin and Auksztulewicz, 2025***). The study showed that while predictive (minimize future surprise) and contrastive (minimize response to expected, maximize to unexpected inputs) PC-inspired learning objectives, implemented using local or global learning mechanisms, exhibited the expected PC-like learning signatures, only the locally trained predictive objective reproduced these signatures more closely than the supervised model (***Gütlin and Auksztulewicz, 2025***). As such, these simulations not only highlight the importance of isolating the specific contribution of learning mechanisms, but also demonstrate that the choice of learning objective and learning mechanism can substantially shape the resulting learning dynamics. This suggests that the potential of PC-inspired learning as a biologically motivated alternative might be closely tied to its specific algorithmic implementation. However, it remains unknown whether predictive or contrastive objectives provide a better empirical account of learning-related neural representations in the brain, and whether local versus global learning mechanisms shape model–brain correspondence over the course of learning.

Here, we address these questions by systematically comparing PC-inspired and supervised learning algorithms within identical recurrent neural network architectures following a controlled modeling approach similar to that used in previous work (***Gütlin and Auksztulewicz, 2025***). Crucially, the current study moves beyond simulations to directly examine the correspondence between artificial and human neural representations during learning. To this end, we compared architecturally identical DNN models using either PC-inspired learning with a predictive or contrastive objective and a local or global learning mechanism, or a conventional supervised objective. We adopted a predictive objective inspired by that of PredNet (***Lotter et al., 2016***), but within an RNN architecture, in which the network learned to predict the next image from preceding inputs. Our contrastive objective was derived from the Forward-Forward algorithm ((***Hinton, 2022***)), in which the network learned to distinguish expected from unexpected inputs. For models with predictive and contrastive objectives, we additionally varied the learning mechanism, applying either local learning, where learning signals were restricted to individual layers, or global learning, where learning signals were propagated across the entire network. Finally, the supervised objective followed a conventional classification approach, in which the network learned to classify the presented image category.

We trained all models on the same statistical learning task used with human participants in a previous EEG study (***McDermott et al., 2026***), allowing us to directly compare model representations with human neural representations based on the same learning task. Specifically, in this task, scene images were presented in pairs, where a leading image from one category probabilistically predicted a trailing image from another category (see ***Figure 1***, left panel). To test whether models using PC-inspired learning algorithms better capture the emergence of learning-related neural representations than conventional supervised algorithms, we used representational similarity analysis (RSA), which allowed us to compare the representational geometry exhibited by each of our models and that observed in the EEG data. Finally, we adopted RSA separately for early and later stages of the original experiment to assess how model–brain correspondence changed over the course of learning.

**Figure 1.**
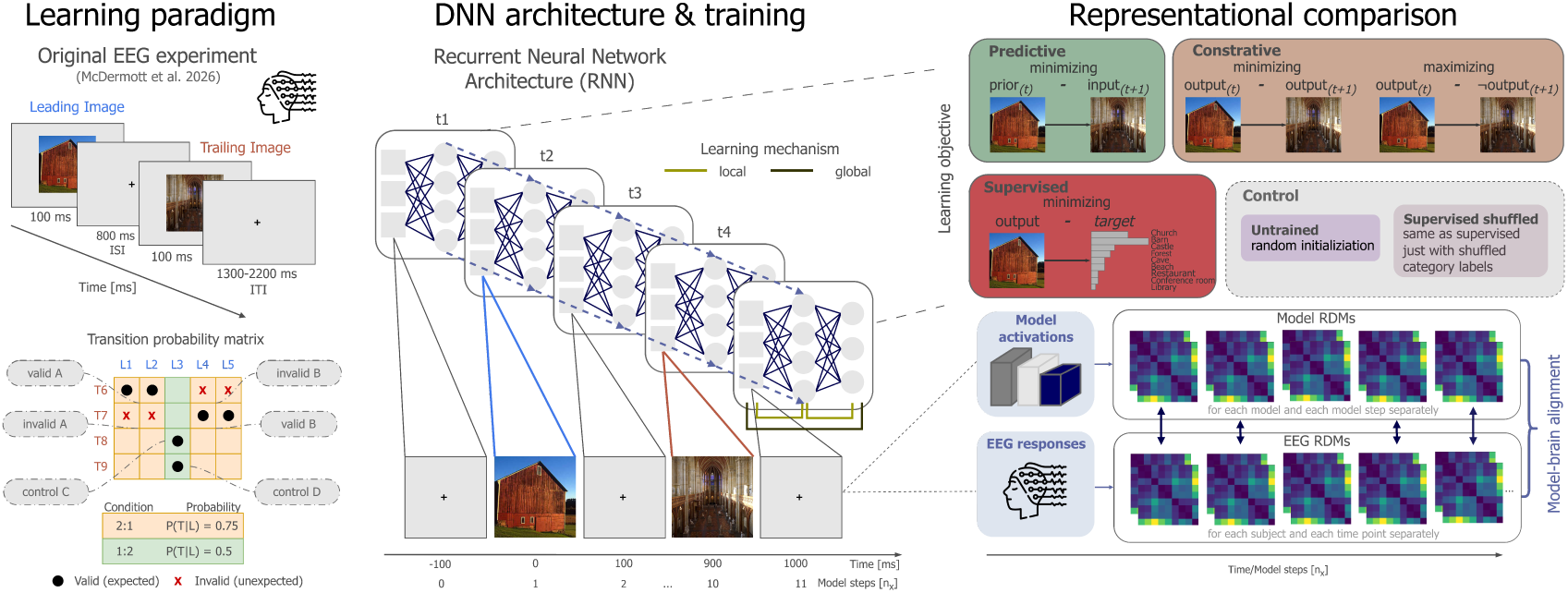
Overview of the modeling approach and model–brain representational comparison. **Left.** Example trial from the statistical learning paradigm used in a previous human EEG study (***McDermott et al., 2026***), in which participants learned associations between leading (blue; e.g., barn) and trailing (red; e.g., church) image categories. Different transition probabilities between leading and trailing image categories were used to facilitate statistical learning, resulting in valid (high probability, 75%), invalid (low probability, 25%), and control (equal probability, 50%) image category pairs (bottom left; figure adapted from McDermott et al. (2026) with permission). For the subsequent representational similarity analysis (RSA), image category pairs were additionally grouped based on trailing image category and transition validity, yielding six category groups: valid A, invalid A, valid B, invalid B, control C, and control D (bottom left, grey). Valid A, for instance, refers to a church image following a leading image category with a 75% transition probability. Center. All models shared the same small recurrent neural network (RNN) architecture, with an image input fed to the network at each successive model processing step (*t*_1_, *t*_2_, …). During training, the models were presented with image sequences mirroring the temporal structure of the statistical learning paradigm used in the EEG study, with one model step per image presentation (100 ms in the EEG paradigm) and eight steps for the inter-stimulus interval (800 ms in the EEG paradigm). Different learning objectives (predictive, contrastive, and supervised) and learning mechanisms (local versus global) were implemented to examine model–brain alignment across these conditions (top right). Both local and global learning were implemented for predictive and contrastive models, whereas the supervised model used global learning only. An untrained model and a shuffled supervised model were additionally included as baseline and control conditions, respectively. **Right.** Neural activity patterns from the EEG data and model activations associated with leading–trailing image pairs were extracted. EEG representational dissimilarity matrices (RDMs) were computed for each participant at each EEG time point, reflecting the pairwise dissimilarities between neural activity patterns associated with the six previously defined category groups (bottom left). Similarly, model RDMs were computed separately for each model across processing steps using the same category groups. Model and EEG RDMs were then compared across model processing steps and EEG time points to assess model–brain representational alignment using RSA.

## Results

### Which optimization algorithm fits brain data best before and after learning?

To determine whether representations learned by PC-inspired models are more closely aligned with neural representations than those learned by supervised models across learning, we used representational similarity analysis in a time-resolved manner (RSA; ***Kriegeskorte et al., 2008***). Central to this approach is the comparison of representational dissimilarity matrices (RDMs), with each RDM quantifying pairwise differences between stimuli or experimental conditions based on their underlying features, thereby allowing representations derived from, for example, computational models and neural data to be directly compared (***Kriegeskorte et al., 2008***). Here, we applied this approach by constructing neural RDMs from EEG data collected during a statistical learning paradigm used in a previous study (***McDermott et al., 2026***). Specifically, we computed RDMs for each each subject and EEG time point relative to stimulus onset using correlation distance (1 − *correlation*), where each RDM reflected pairwise dissimilarities between EEG responses averaged across trials sharing the same trailing image category and transition validity (valid, invalid, or control; see ***Figure 1***, bottom left panel). In other words, each EEG RDM captured neural representations for specific combinations of trailing image category and transition validity, such as valid church and invalid church (for more details, see EEG RDM extraction). We applied the same representational approach to each of our models, which were trained on the same statistical learning task as that used in the EEG study (for more details, see Experimental stimuli and training). Hence, we computed model RDMs at each model processing step for our four PC-inspired models resulting from the combinations of learning objective and learning mechanism (i.e., Contrastive global, Contrastive local, Predictive global, and Predictive local), as well as for the Supervised model (Figure 1, center and right panels), with pairwise dissimilarities capturing differences in model representations across the same category groups as in the EEG-derived RDMs (Model conditions). As we additionally included two control models, corresponding RDMs were also computed for the Supervised shuffled model to assess the effect of training without meaningful category labels, and the Untrained model to assess the contribution of network architecture alone. Lastly, to quantify the representational alignment between neural and model RDMs, we calculated Spearman’s rho, where stronger positive correlations would indicate greater similarity between the two RDMs and, consequently, greater representational alignment between neural and model representations. To determine which model overall best accounted for the observed neural data during early (first half) and late (second half) stages of learning, we used Bayesian Model Selection to compare model–brain correspondence separately for the first and second half of the experiment using the Spearman’s rho coefficients obtained from the EEG-model RDM comparisons. Given that our time-resolved RSA approach compared EEG and model RDMs across multiple EEG time points and model steps for each subject, these comparisons yielded multiple Spearman’s rho coefficients. We therefore averaged, for each subject, the Spearman’s rho coefficients across EEG time points and model steps following the onset of the trailing image. These subject-level averages were then used for Bayesian Model Selection (see Model comparison).

Bayesian Model Selection revealed that during the first half of learning, neural representations were best captured by the Supervised model (*P* (best) = .932, corresponding to strong evidence over alternative models), whereas during the second half of learning, the Predictive local model showed the strongest representational correspondence with the observed neural representations (*P* (best) = .769, corresponding to moderate evidence over alternative models). Besides these two models, all remaining models failed to provide a comparable account of the neural representations (***Figure 2***). To formally test this observed pattern, we fitted a Hierarchical Bayesian model investigating the interaction over model type (Supervised vs. Predictive local) and learning stages (first vs. second half). Consistent with the earlier observed pattern, we found moderate evidence for the interaction between model and the first and second half of learning (*BF*_10_ = 6.00; effect mean = .032, 95% HDI [−.029, .089], *P* (*θ* > 0) = .857), indicating an interaction between the process of learning and the emergence of brain representations associated with predictive rather than category-based processes. Together, these findings indicate two patterns: First, the Supervised model and, among all PC-inspired models, only the Predictive local model exhibited close representational alignment with the neural representations derived from EEG data. Second, the relative correspondence between model and neural representations varied across learning stages, with neural representations becoming increasingly aligned with predictive representations over the course of learning.

**Figure 2.**
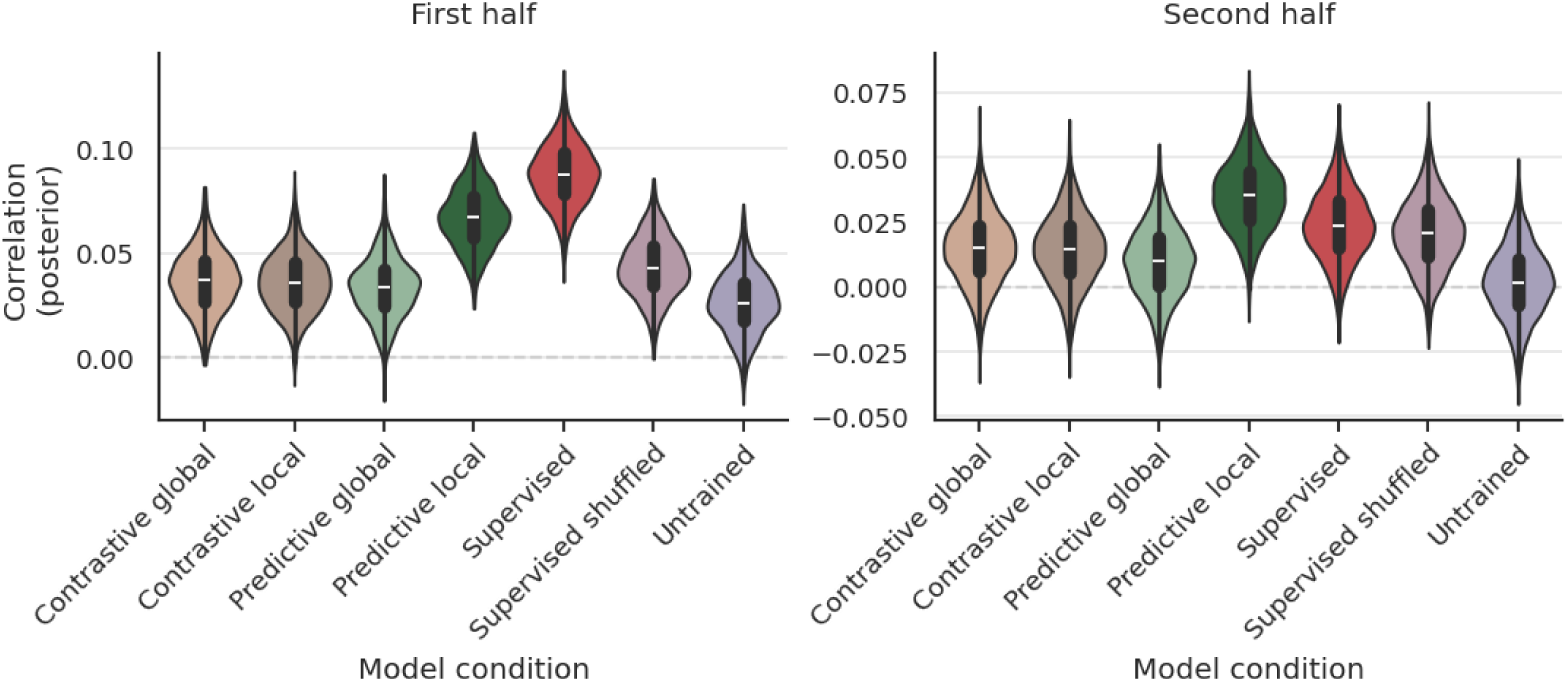
Representational alignment between brain responses and model representations shifts across learning. Violin plots show the posterior distributions of the estimated mean model–brain representational similarity for each model condition, based on Spearman correlations between model and EEG RDMs during the first (left) and second (right) halves of the experiment. Bayesian model selection (BMS) estimated the posterior probability (*P* (best)) that a given model has the highest mean representational similarity among the compared models, based on these posterior distributions. **Left.** During the first half, the Supervised model showed the strongest representational similarity to neural representations derived from EEG data. **Right.** During the second half, the Predictive local model showed the highest estimated mean representational similarity, while similarity for the Supervised model declined.

### Does the Predictive local model explain unique information about the brain?

To determine which aspects of neural representations after learning were uniquely captured by the Predictive local model, we performed a time-resolved residual correlation analysis to examine its unique correspondence with EEG RDMs during the second half of the experiment (i.e., after learning). To isolate the correspondence specifically associated with the Predictive local model, we first removed variance in the Predictive local RDMs explained by the Supervised (as the main alternative), as well as the Untrained and Supervised shuffled models (as controls). Accounting for variance associated with these models then allowed us to focus on unique model-brain correspondence beyond that explained by model architecture (i.e.,through the Untrained model) or category-based information (i.e.,through the Supervised models). Finally, to assess the correspondence of this unique variance with neural representations, we compared the resulting residual RDMs with the EEG RDMs using Spearman’s rho at each model step and EEG time point.

Following FDR correction, our analysis revealed several significant clusters of correspondence across the intersections of leading- and trailing-image representations in the model and EEG data (***Figure 3***). Specifically, correspondence was particularly pronounced between model representations following leading-image onset (model steps 1-9) and EEG leading-image representations (0-0.5 s), indicating that the Predictive local model may form leading image category representations that resemble those observed in the EEG data but are not present in the Untrained, Supervised, and Supervised shuffled models. In contrast, correspondence between model representations following trailing-image onset (model steps 10-14) and EEG trailing-image representations (0.8-1.4 s) was comparatively weaker. More interestingly, the analysis additionally revealed a notable correspondence between model representations following leading-image onset (model steps 1-9) and EEG trailing-image representations (0.8-1.4 s) and vice versa (model trailing-image and EEG leading-image), suggesting that the Predictive local model may not simply form representations confined to the individual image categories, but instead integrates information across leading- and trailing- image categories, reflecting its predictive learning objective. Thus, the closer representational alignment between Predictive local and neural representations during the second half of learning may arise, at least in part, from the model’s ability to form representations that extend beyond the currently presented stimulus category.

**Figure 3.**
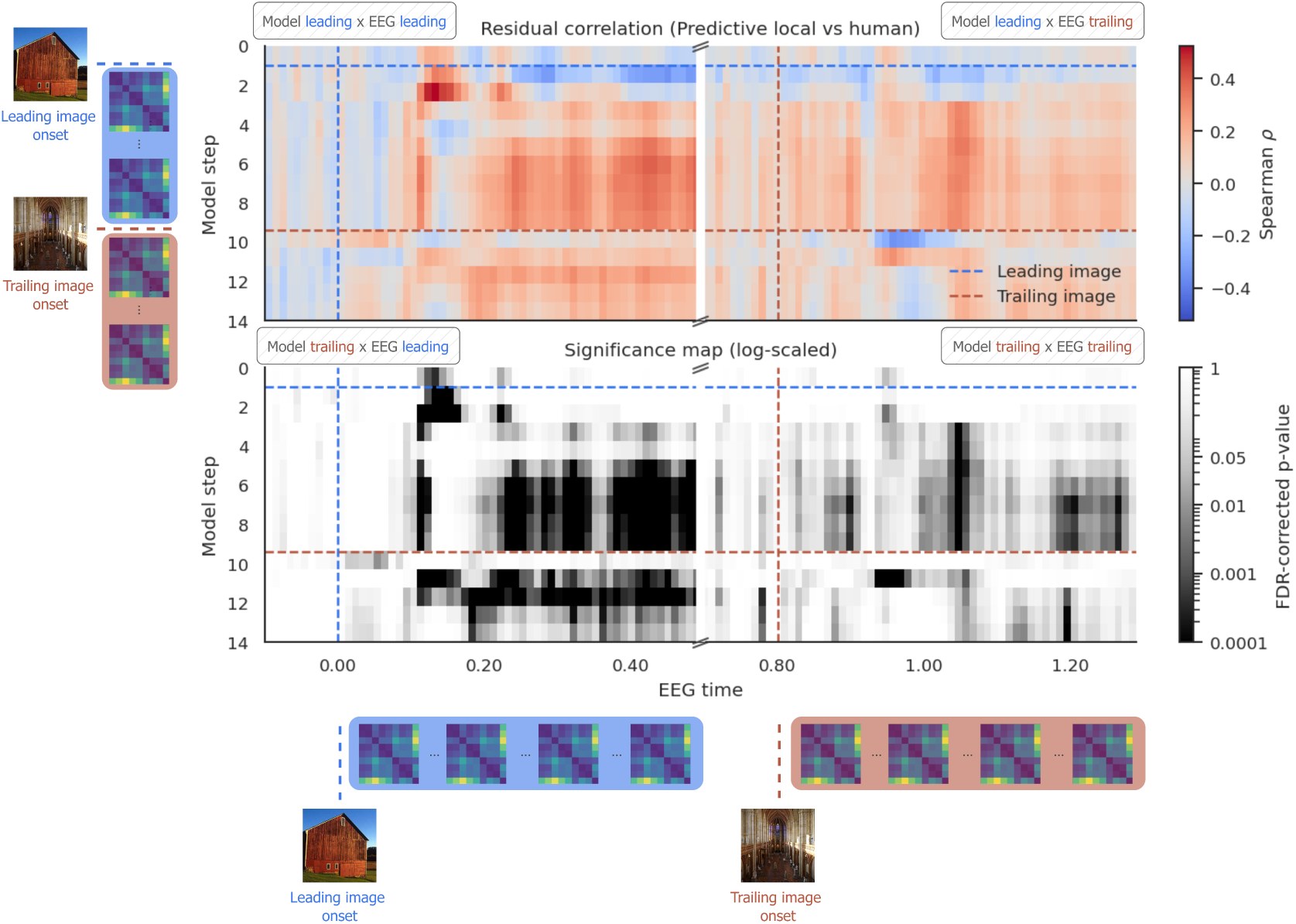
Residual representational alignment between the Predictive local model and the brain after controlling for other model conditions. The colormesh map displays correlations between EEG RDMs and residual RDMs of the Predictive local model after removing variance explained by the Supervised, Supervised shuffled, and Untrained model conditions, across EEG time points and model processing steps (top). Red shades indicate positive correlations, with darker red indicating stronger positive correlations, whereas blue shades indicate negative correlations, with darker blue indicating stronger negative correlations. The grayscale map displays the statistical significance of the corresponding correlations, with darker shades indicating lower FDR-corrected p-values (bottom; *r* > 0, FDR-corrected, *α* = .01). Onset times of the leading and trailing images are indicated by blue and red dashed lines, respectively, along both the EEG time and model steps. An example leading–trailing image sequence is shown next to the model step axis and below the EEG time axis to illustrate a stimulus sequence underlying the comparison of model and EEG representations over time. Significant correlations are observed across the entire map, with particularly pronounced clusters at the intersections of model leading-image and EEG leading-image representations, model trailing-image and EEG leading-image representations, and model leading-image and EEG trailing-image representations. In contrast, the intersection of model trailing-image and EEG trailing-image representations shows comparatively weaker correlations, although some significant clusters remain.

### What type of information does the Predictive local model capture?

To characterize the specific information shared between Predictive local and EEG data, we compared those representations against a set of theoretically motivated templates using RSA. To this end, we constructed three template RDMs: a *stimulus-driven RDM*, reflecting the identity of the presented trailing stimulus category irrespective of prediction validity (i.e., valid vs. invalid image pairs); a *prediction-driven RDM*, reflecting the predicted stimulus category irrespective of the actual presented stimulus category; and a *validity-driven RDM*, reflecting sensitivity to prediction validity (valid vs. invalid image pairs), irrespective of stimulus identity or predicted category (see ***Figure 4*** bottom right).

**Figure 4.**
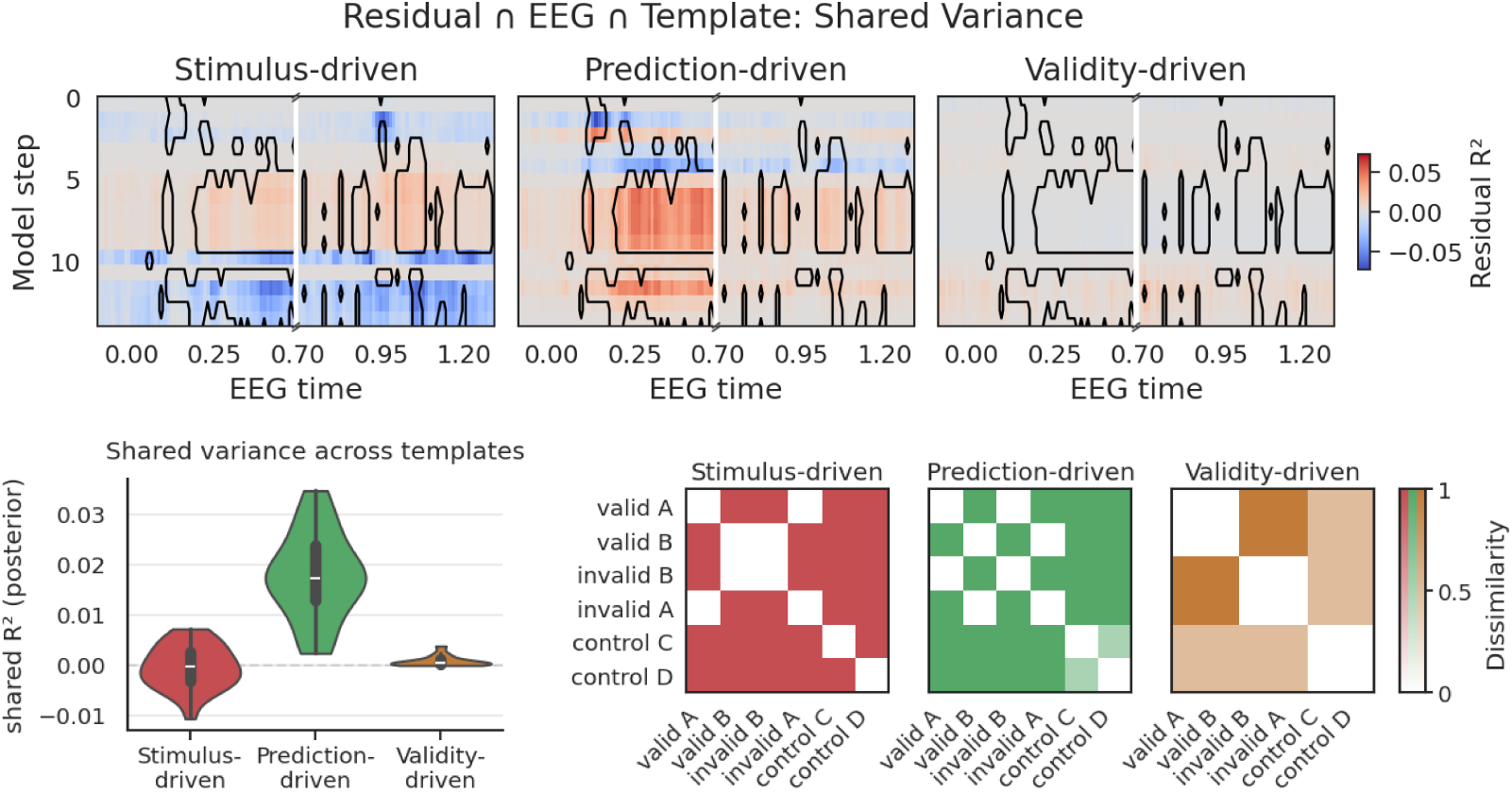
Shared variance between the Predictive local model and the brain primarily reflects prediction-driven information. Three template RDMs (stimulus-driven, prediction-driven, and validity-driven information; bottom right) were used to characterize the shared variance between the Predictive local model and EEG data across the first and second halves of the experiment. **Top.** Shared variance between EEG data, the Predictive local model, and each template across model steps and EEG time points, shown within the significant clusters identified in the preceding analysis (see Figure 3), which are outlined in black. The prediction-driven template captured the largest amount of variance shared between the Predictive local model and EEG data, with shared variance already evident immediately after the model had processed the leading image (top center). The stimulus-driven template showed opposing patterns of shared variance during leading- and trailing-image processing, resulting in a near-zero mean (top left), whereas the validity-driven template showed only a small amount of shared variance after presentation of the trailing image (top right). **Bottom left.** Mean shared variance between the Predictive local model and EEG data for each template, averaged across model steps and EEG time points within the significant clusters, with the prediction-driven template showing the highest mean shared variance.

Comparing the three template RDMs against the residual Predictive local-EEG correspondence revealed that the underlying representations were best explained by the prediction-driven template, whereas substantially weaker effects were observed for the stimulus-driven and validity-driven templates (***Figure 4***, bottom left).

Inspection of the heatmaps (see ***Figure 4*** top) showed that, for the prediction-driven template, shared variance was most pronounced between model leading-image and EEG leading-image representations, as well as between model trailing-image and leading-image EEG representations. Weaker but still positive correspondence was observed between the model leading-image and EEG trailing-image representations. For the stimulus-driven template, positive correspondence between model leading-image and EEG leading-image representations was accompanied by negative correspondence between model trailing-image and EEG leading-image representations, suggesting that these effects partially canceled each other out and thereby contributed to the comparatively weaker overall correspondence observed for the stimulus-driven template. The validity- driven template showed little to no shared variance across all intersections.

In line with the patterns observed in the heatmaps, averaging across all significant clusters showed substantially greater shared variance for the prediction-driven template (mean = 0.018, 95% HDI [0.015, 0.021], *P* (*θ* > 0) = 1.000), while shared variance was close to zero for both the stimulus-driven template (mean = 0.000, 95% HDI [−0.002, 0.001], *P* (*θ* > 0) = .304) and the validity-driven template (mean = 0.001, 95% HDI [0.000, 0.001], *P* (*θ* > 0) = 1.000) (see ***Figure 4*** bottom left). Together, these results indicate that the unique shared variance between the Predictive local model and brain representations primarily reflects predicted stimulus categories rather than presented stimulus categories or prediction validity (general mismatch signals).

## Discussion

In this study, we investigated to what extent different PC-inspired DNNs capture brain representations compared to supervised DNNs across learning by systematically varying both the learning objective (contrastive versus predictive) and learning mechanism (local versus global) of PC-inspired DNNs. We trained architecturally identical RNNs on a statistical learning task and compared their representations to human EEG data (***McDermott et al., 2026***) recorded during the same paradigm using RSA. We show that given otherwise equal conditions, only the Predictive local model learns representations that are more closely aligned with human brain representations than those of the supervised DNN, and that this effect emerges only after learning. Crucially, this suggests that closer model-brain alignment does not arise from predictive-coding-inspired learning algorithms per se, but depends on their specific implementation, including the learning objective and learning mechanism. More broadly, this aligns with recent work emphasizing the importance of learning objectives and learning dynamics in shaping representational alignment between DNNs and the brain (***Raugel et al., 2026***). Our results further highlight prediction as a key component underlying closer model-brain alignment during learning.

### Not all PC-inspired models resemble brain representation better than supervised models

It is a longstanding question whether PC-inspired DNNs can explain brain representations in a way that supervised DNNs cannot (***Brands et al., 2025***; ***Fonseca, 2019***; ***Leutenegger, 2026***; ***Rane et al., 2020***). Our findings add evidence in support of this by showing that the Predictive local model outperformed the supervised model after learning. However, our results also indicate that this advantage is not shared across all PC-inspired models, and may depend on the specific PC-inspired learning algorithm implemented. This result is particularly relevant as existing PC-inspired models as such constitute a family of computational models that implement predictive coding principles in different ways (for an overview see ***Millidge et al., 2021***). For example, some PC-inspired models primarily employ local learning mechanisms but optimize a supervised objective, i.e., learning to predict specific labels or observations (e.g., ***Whittington and Bogacz, 2017***; ***Tscshantz et al., 2023***). Others instead optimize a predictive objective, i.e., learning to predict future inputs, while relying on global learning mechanisms such as backpropagation (***Lotter et al., 2016***); or combine both predictive learning objectives with local learning mechanisms (***Gütlin and Auksztulewicz, 2025***; ***Millidge et al., 2024***). Our findings suggest that both the learning objective and the learning mechanisms are functionally relevant for the alignment between PC-inspired models and the brain, since only the PC-inspired model that combined a predictive objective with local learning mechanisms formed representations that showed significant similarity to the brain.

Despite previous work demonstrating that contrastive learning (which can be PC-compatible; see ***van den Oord et al., 2018***; ***Hinton, 2022***) can show similar or better brain alignment than supervised learning (***Conwell et al., 2024***), we found no evidence for such an advantage in the present study. One possible explanation is that predictive and contrastive objectives favor different representational formats due to their distinct learning objectives, and that these representations differ in their relevance for the statistical learning task used here. That is, in the current task, the predictive objective may promote learning representations capturing relationships between successive stimuli, whereas the contrastive objective may promote learning representations that distinguish between expected vs. unexpected pairs. In line with the proposals that statistical learning results in the formation of stimulus-specific associations rather than abstract rules (***Turk-Browne et al., 2009***), predictive objectives may therefore be particularly well suited to capture the representations emerging during statistical learning, which potentially explains why only the Predictive local model showed closer alignment with the brain. It should be noted that our conclusion is limited to the specific formulation of the contrastive objective used here. Motivated by its explicit PC interpretation, we employed the contrastive objective proposed in the Forward-Forward algorithm (***Hinton, 2022***), which encourages the network to distinguish positive input data (here expected pairs) from negative input data (here unexpected pairs). The Forward-Forward algorithm has the unique attribute that the contrast between expected and unexpected is only evaluated at the level of overall layer activity (i.e. minimizing/maximizing the distance of vector norms rather than the full vector for positive/negative samples). This is different from other formulations of contrastive objectives, such as Contrastive Predictive Coding (CPC; ***van den Oord et al., 2018***), where contrasting different inputs happens within the same node (i.e., the unreduced latent vector), potentially resulting in different learned representations. Future work should therefore investigate whether alternative contrastive objectives, such as CPC, yield representations that align more closely with brain representations during statistical learning.

### Learning-stage-dependent correspondence between model and brain representations

A key question raised by our findings is why the supervised model aligned more closely with brain representations than the Predictive local model prior to learning. One possible explanation may lie in the dynamics of statistical learning in the brain itself. There is vast evidence that brains perform high level object recognition (***DiCarlo and Cox, 2007***; ***Peelen and Downing, 2017***; ***Grill-Spector and Weiner, 2014***). At the same time, statistical learning has been shown to progressively reshape brain representations over the course of learning as regularities and associations between stimuli are acquired, such that representations increasingly reflect the regularities of the environment by increasing the representational similarity between associated stimuli (***Greco et al., 2024***). Such learning-dependent changes in neural representations may therefore differentially favor representations which map onto the different learning objectives of the models, with the supervised model primarily learning to discriminate between categories, whereas the Predictive local model explicitly learns to predict upcoming inputs and therefore may be particularly sensitive to the statistical relationships between successive stimuli. From this perspective, the supervised model may better capture brain representations at earlier stages of learning, when category information is based on prior semantic knowledge but the statistical relationships between stimuli have not yet been acquired. In contrast, as statistical learning unfolds and neural representations increasingly reflect the learned structure of the environment, the Predictive local model may better capture the predictive relationships between stimuli and therefore show closer alignment with neural representations after learning.

### Unique variance captured by the Predictive local model

Previous work has suggested that PC networks may capture representational information complementary to that learned by supervised DNNs (***Fonseca, 2019***). Our current findings extend this account, showing the Predictive local model explained variance in brain representations beyond that accounted for by the supervised model. Importantly, the unique Predictive local–brain alignment was not restricted to representations of matching image categories (i.e., leading–leading and trailing–trailing), but extended across representations of leading and trailing image categories. In other words, representations of leading stimuli appeared to contain information related to trailing stimuli (Model leading X EEG trailing) and vice versa (Model trailing x EEG leading), reflecting a pattern that would be expected given the predictive learning objective of the Predictive local model. One interpretation of this pattern is that the complementary information captured by the Predictive local model beyond the supervised DNN may relate to relationships between successive stimuli rather than stimulus-category information alone. Such relational representations have been repeatedly reported in the statistical learning literature, where representations of associated stimuli have been shown to become more similar following the exposure to statistical regularities. In this context, some studies demonstrated that statistical learning can shape representations in a bidirectional associative manner, whereby the representation of one stimulus becomes increasingly similar to the representation of its associated stimulus and vice versa (***Schapiro et al., 2012***), or in a predictive (forward) looking manner, whereby representations of preceding stimuli become increasingly shaped by information about subsequent stimuli, reflecting the temporal structure of the learned sequence (***Schapiro et al., 2012***; ***Greco et al., 2024***). More recent accounts, however, suggest that predictive representations acquired through statistical learning may not be restricted to forward-looking relationships, but may additionally encode information about preceding stimuli (***Sharp and Eldar, 2024***; ***Tummeltshammer et al., 2017***). This may also explain why we observed significant correspondence not only between model leading-image and EEG trailing-image representations, but also between model trailing-image and EEG leading-image representations. Taken together, our findings not only provide insight into the nature of the complementary information captured by PC-inspired models relative to supervised DNNs, but also suggest that the representational patterns learned by the Predictive local model resemble those reported for neural representations following statistical learning.

### Predictive local representations reflect predicted not presented stimulus information nor prediction validity information

Beyond demonstrating that the Predictive local model captures information stimulus-category relationships, our findings further demonstrated that the shared representations between the Predictive local model and the brain were specifically driven by expected stimulus category rather than the presented stimulus category or prediction validity. Unlike our previous analyses, which primarily informed where the unique correspondence between the Predictive local model and the brain emerged, the present findings provide insight into the nature of the information underlying this correspondence. Our finding is consistent with accounts proposing that prediction plays a central role in (statistical) learning, enabling the extraction of regularities by anticipating future events based on past experience (***Friston, 2005***; ***De Lange and Press, 2026***).

### Future directions

Our results demonstrate that predictive mechanisms provide a better model of the brain than contrastive mechanisms, given otherwise fixed circumstances. However, the literature indicates that contrastive methods are highly effective for learning (***Chen et al., 2020***) and can perform well on brain modelling tasks (***Conwell et al., 2024***). Accordingly, it may be valuable to investigate other contrastive algorithms in future work. Beyond this variation in algorithms, it remains to be investigated whether these effects reproduce across different scales and architectures of neural networks, and it would be worthwhile for future studies to examine how these algorithms perform at larger scale and with other variants. When considering the scaling of these networks, it is important to note that a general limitation of PCNs is that, to date, no strictly PCN-aligned model has been shown to scale to the same level of abstraction achieved by classical deep learning models. This issue of scaling PCNs to match other commonly used DNNs is addressed in a separate forthcoming publication (***Gütlin et al., 2026***).

### Summary

In this study, we investigate predictive-coding-inspired neural networks and isolate the effect of the learning algorithm by training otherwise identical networks on a statistical learning task. We find that a Predictive local algorithm shows the strongest correspondence to the brain after learning, and that this model uniquely captures variance in brain data that cannot be attributed to its architecture or other factors. We further show that the unique variance explained by the Predictive local model is more consistent with predictive signals than with category-specific or validity-based (mismatch) signals. These findings suggest that the brain progressively attenuates category information while retaining and refining predictive structure, underscoring the critical role of predictive processing in shaping neural representations over the course of learning.

## Methods and Materials

### Model setup

To isolate the effects of learning algorithms, we employed strictly parameter-matched models across all conditions. Instead of comparing DNNs with different architectures and numbers of parameters to maximize performance, we matched model architectures and capacity across all conditions to maximize interpretability. This design ensures that any differences in model behavior and representations can be directly attributed to the optimization procedure rather than to architectural or capacity differences. We used Simple RNN architectures with hyperparameters matched across all conditions except for the learning objective and learning mechanism (local vs. global). Each model consisted of two Keras Simple RNN layers (one feedforward kernel and one recurrent kernel per layer) with 256 and 128 latent units respectively (1,343,872 trainable parameters overall), leaky ReLU activation and Keras default parameters for all unspecified settings.

### Model conditions

Models were trained under five learning objective conditions and two learning mechanism conditions (see ***Figure 1***, center and top-right panels), resulting in a flexible factorial manipulation of the training procedure with a total of seven conditions outlined below.

### Learning Objective

The main manipulation applied in this study is the learning objective, defined as the target state toward which the network is instructed to converge via the loss function. Five conditions were tested: *Contrastive*, *Predictive*, *Supervised*, *Supervised shufled*, *Untrained*.

In the Contrastive condition, the network was trained following the objective proposed by Hinton (2022), applied to a standard recurrent neural network architecture, minimizing the summed squared layer activation for valid pairs while maximizing it for invalid pairs. This algorithm is conceptually related to predictive coding (PC), whereby expected (valid) stimuli evoke minimal activity and unexpected (invalid) stimuli evoke maximal activity.

In the Predictive condition, the model learned by predicting the next frame from prior frames. The loss was defined as the mean squared error (MSE) between the predicted future frame, obtained by projecting the prior latent state (the latent frame multiplied by the recurrent kernel) back onto the input space, and the true future frame. Accordingly, the model learns a latent representation of the input over time. This implementation is closely related to autoregressive video prediction models such as Assran et al. (2025), and more specifically to Lotter et al. (2016), in which the learning signal arises from an autoregressive prediction of the future state by minimizing the difference between the latent state and an explicitly modeled error state within a recurrent neural network. Unlike PredNet, which relies on convolutional and convolutional LSTM layers, the error state in the present model is represented within a simple recurrent neural network.

In the Supervised condition, the learning objective followed a standard supervised classification scheme. At each timestep, the objective was to correctly classify the presented stimulus category using a categorical cross-entropy loss (see ***Krizhevsky et al., 2012***; ***Hastie et al., 2001***). For this purpose, a dedicated readout layer was added, projecting the output of the last recurrent layer onto a one-hot logit encoding for each classification category.

We introduced the Supervised shuffled condition as a control for the Supervised condition, which is identical except that the correct labels were randomly shuffled prior to fitting, thereby eliminating the learning signal by rendering the target effectively meaningless. In contrast to the Untrained condition, the weights in this condition were still updated during training (which may itself induce brain-like structure), but on a meaningless target.

As an additional baseline, the Untrained condition used a network with the same architecture as all other conditions, but without any training. This condition indicates the extent to which brain similarity can be attributed to the network architecture itself rather than to the training process.

### Learning Mechanism

In addition to the learning objective, the predictive and contrastive algorithms allow us to implement a version of the network that does not rely on full backpropagation loops. This means that for these learning objectives we can manipulate the learning mechanism (the operational step by which the network is instructed to reach the learning objective) by using either full global backpropagation gradients or partial, localized gradients. In the Global condition, the loss is fixed at the final layer, and all weights are updated using a full, standard backpropagation loop throughout the entire network. In the Local condition, gradient propagation is restricted to a single layer at a time: each layer’s output is trained against a learning objective defined solely with respect to the output of the previous layer, such that layer N predicts or contrasts based on the output of layer *N* −1, with the original input used only as input to the first layer. This results in a more localized credit assignment across the network. Taken together, the model space included seven models: Contrastive global, Contrastive local, Predictive global, Predictive local, Supervised, Supervised shuffled, and Untrained.

### Experimental stimuli and training

To isolate the effect of learning, we trained our DNN models under the same experimental setup as the subjects in the original study (***McDermott et al., 2026***). The procedure is described below.

### Experimental setup

We used a dataset from a study by McDermott et al. (2026), which aimed to investigate the underlying mechanisms of expectation suppression during statistical learning. This dataset includes EEG recordings from 31 participants (25 female, 5 male, 1 non-binary) between 18 and 35 years (*mean age* = 22.7 years), as well as the image material used during the task. EEG data were acquired at a sampling rate of 2048 Hz using a BioSemi ActiveTwo system with 64 electrodes arranged according to the international 10/20 system, recorded while participants performed a statistical learning task. Preprocessing of the EEG data followed a standard protocol, including high-pass filtering to remove low frequencies below 0.1 Hz, notch-filtering between 48-52 Hz using a zero-phase Butterworth filter, downsampling to 300 Hz, re-referencing to average and epoching from -200 to 2600 ms relative to stimulus onset. Additionally, baseline correction to the prestimulus period and low-pass filtering of 48 Hz were applied. The image material in the study by McDermott et al. (2026) consists of 3 586 color images of natural scenes representing nine distinct categories. Given its combination of EEG data on a learning task and a relatively large amount of image material for a neuroimaging study, this dataset provided a comprehensive foundation for investigating the correspondence between learning in the brain and biologically inspired DNNs.

### Experimental paradigm

During the EEG recording, participants performed a statistical learning task in which they learned the transitional probability between different image categories, i.e., the probability of one image category predicting images from another category. In each trial, participants were first presented with a leading image, randomly selected from five possible leading categories (*barn*, *beach*, *library*, *restaurant*, *cave*), followed by a trailing image from one of four possible trailing categories (*church*, *conference room*, *castle*, *forest*). This structure allowed participants to make a probabilistic prediction about the trailing image (category), which could result in either a valid prediction (expected image category pair, occurring at 75% chance), a invalid prediction (unexpected image category pair, occurring at 25% chance). Additionally, there was a control condition, where each leading category was followed by either of 2 trailing categories with 50% chance. Each of the images were only shown once to avoid repetition suppression effects. Overall, participants completed eight blocks of 216 trials each without any prior training. Throughout the task, trial order was randomized. To maintain participants’ attention throughout the task, 5% of the trials featured images that were displayed upside down, prompting participants to press a button whenever these images appeared. The full trial structure and transitional probability are illustrated in the left panel of ***Figure 1***.

### Model inputs and training

Model inputs were constructed to exactly match the experimental paradigm (see Experimental paradigm). The model was fed the same sequence of images as human subjects, so that each input sequence corresponded to the stimuli presented during one trial, with 100 ms of human subject time mapping onto one step in the model sequence. Specifically, each sequence consisted of 1 step gray mask, 1 step leading image, 8 steps mask (inter-stimulus interval), 1 step trailing image, and 4 steps gray mask. The stimulus data were derived from the material of the original study and further extended with additional images created for each category. 800 samples (80 per leading category and 100 per trailing category) from the original images were separated to serve as the test set and be used for the analysis, while the rest of the original images plus the additional images were used for training (for a total of 6392 stimuli and an average of 710 samples per category). For network input, images were resized to 40×40 pixels, normalized to the range [0, 1], and flattened across the height, width, and channel dimensions into a single feature vector, which was then fed into the RNN architecture. These image sequences were used to train all seven models (see Model conditions) for 100 epochs, with 3196 random combinations drawn per epoch from the stimulus material. For each model condition, the epoch with the best validation loss across the 100 training epochs was selected for further analysis. As optimizer, Adam was used with standard settings.

### Representational Similarity Analysis

The representational similarity between models and the brain was compared using representational similarity analysis (RSA; ***Kriegeskorte et al., 2008***). The representational dissimilarity matrices (RDMs) were constructed based on a combination of the trailing image category (simplified by letter codes: *church = A*, *conference room = B*, *castle = C*, *forest = D*) and whether the trailing image was preceded by a *valid* (75% probability), *invalid* (25% probability), or *control* (50% probability) leading category (see Experimental paradigm). For example, a church image following a leading image with a transition probability of 75% was considered part of the *valid church* (i.e., valid A) category group, and likewise for *invalid church* (i.e., invalid A) with a transition probability of 25%. This categorization captured both category-level and statistical-learning validity information.

### Model RDM extraction

To extract RDMs from the models, images were drawn from the test dataset according to the category mapping defined in Representational Similarity Analysis, and preprocessed as described in Model inputs and training. The models were then evaluated on each category, and the neural activation patterns of the latent states of both RNN layers were extracted and concatenated. Activation patterns were averaged across samples within each category and pairwise distances between category means were computed over the activation vector using correlational distance (1 − *correlation*).

### EEG RDM extraction

Epoched EEG data from the original study were loaded (for preprocessing details, see ***McDermott et al., 2026***) and resampled to 100 Hz using MNE-Python (***Larson et al., 2023***). Category groups were defined based on the above described mapping (see Representational Similarity Analysis). EEG trials were extracted according to these category groups and split into two halves representing the first 50% and last 50% of trials to investigate brain representations at early (first half of experiment) vs. late stages (second half of the experiment) of statistical learning. Trial averages were computed across all trials within each category group. For each subject and each timestep, pairwise distances (via correlation distance: 1 − *correlation*) were calculated between category groups using EEG channels as features, yielding one RDM per timestep and subject.

### Statistical evaluation

Statistical group comparisons were performed using Bayesian regression models implemented in bambi (***Capretto et al., 2020***). RSA methods and residual correlation were implemented using scipy (***Virtanen et al., 2020***).

### Model comparison

To compare the model conditions, model RDM and EEG RDM were correlated using Spearman rank correlation for each combination of DNN model, EEG subject, DNN time step, and EEG time step. This was performed separately for the first and second half of the training data. For the final comparison, the highest average correlation after the onset of the second image was extracted.

For each data split, a Bayesian hierarchical model was fit predicting correlation strength from model condition, with subject-specific random intercepts to account for the paired samples structure. Model comparison was performed via posterior ranking: for each posterior sample, the model with the largest estimated mean correlation was identified, and the proportion of samples in which each model ranked first was calculated. These proportions are reported as the posterior probability that a given model is the best-performing model, *P(best)*.

### Model interaction over experiment

The Supervised and Predictive local models were selected as the best-performing models for further analysis of their correspondence to EEG data across the early and late phases of the experiment. To this end, correlation data from these two models were used to fit a Bayesian regression model examining the interaction between model condition and EEG data split (first vs. second half of experimental trials), with EEG subject included as a random effect.

### Residual correlation

To investigate where Predictive local explains unique variance in the EEG data, a residual correlation approach (see ***Cohen et al., 2013***) was applied for each model time step and EEG time point. For each combination of model step, EEG time, and subject, the variance in the Predictive local condition that is linearly explainable by the Supervised, Supervised shuffled, or Untrained conditions was factored out. To this end, a non-negative least squares (NNLS) regression was fit with Predictive local RDMs as the target and the Supervised, Supervised shuffled, and Untrained RDMs as predictors. The predicted RDM obtained from this regression was then subtracted from the original Predictive local RDM, yielding a residual RDM containing only variance not linearly explainable by the other model conditions.

To assess where the residual correlation across subjects was significantly greater than zero, a one-sample z-test was applied to Fisher z-transformed correlation values (***Fisher, 1915***). Multiple comparisons across model steps and EEG time points were corrected for using false discovery rate (Benjamini–Hochberg FDR with *α* = .01) correction applied to all individual tests.

### Template shared variance

To investigate the nature of the correspondence between the DNN model and EEG data, three template RDMs were constructed, each representing the expected dissimilarity pattern under the assumption that the data were perfectly determined by a specific component:

1.The **stimulus-driven template RDM** describes the pattern that would arise if the data were perfectly determined by the trailing stimulus class.
2.The **prediction-driven template RDM** describes the pattern that would arise if the data were perfectly determined by the stimulus class expected during that trial.
3.The **validity-driven template RDM** describes the pattern that would arise if the data were perfectly determined by whether the trailing image was expected or unexpected.

These template RDMs were then used to calculate the variance jointly shared between the Predictive local model, the EEG data, and each template. To this end, NNLS regression was used to predict the EEG data using the template RDM alone (*R*^2^(EEG ∣ template)), the Predictive local residual RDM alone (*R*^2^(EEG ∣ residual)), and both as stacked predictors (*R*^2^(EEG ∣ residual, template)). The shared variance was calculated as

*R*^2^(EEG ∩ residual ∩ template) = *R*^2^(EEG ∣ residual) + *R*^2^(EEG ∣ template) − *R*^2^(EEG ∣ residual, template).

It should be noted that this method, in common with other shared and unique variance approaches, can yield negative R² values, which typically reflect interactions between predictor variables where the combination of predictors explains more variance than either alone (***Schoen et al., 2011***; ***Mood, 1971***).

The above joint variance procedure procedure yielded one map per template (each of dimensions: model step x EEG time x subject), describing the variance shared between that template, the EEG data, and the DNN model. For each template, indices where the residual correlation reported in Residual correlation was significant (*p* < .05) were selected, and the average shared variance within these regions was compared across subjects using a Bayesian regression, testing whether the correlation for each template significantly deviated from zero. This allowed us to identify which of these representational components likely contributed to the correspondence between the DNN model and the EEG data.

## Acknowledgments

We thank Hannah McDermott for providing access to the stimulus materials and EEG data from her study for use in the present study, and for the helpful discussions regarding her experimental paradigm and findings. Further, we thank Mahdi Enan for comments and suggestions on the manuscript. Finally, we thank Karlo Trupec for reviewing our code. This work was performed using the Curta HPC cluster at Freie Universität Berlin (***Bennett et al., 2020***). The EEG icon in ***Figure 1*** was obtained from Flaticon (https://www.flaticon.com).

## Data availability statement

All code required to replicate this study is available at: https://github.com/PredLabUM/representations_of_predictive_coding_networks. The original data from McDermott et al. (2026) are available at: https://osf.io/x7ydf.

## Additional information

### Author contributions

**Dirk Gütlin:** Conceptualization, Data Curation, Formal Analysis, Investigation, Methodology, Project Administration, Software, Supervision, Validation, Visualization, Writing - Original Draft Preparation, Writing - Review & Editing. **Denise Kittelmann:** Conceptualization, Formal Analysis, Investigation, Methodology, Project Administration, Software, Validation, Visualization, Writing - Original Draft Preparation, Writing - Review & Editing. **Ryszard Auksztulewicz:** Conceptualization, Funding Acquisition, Investigation, Methodology, Project Administration, Resources, Supervision, Validation, Writing - Review & Editing.

### Statement of Authorship and Verification

The authors affirm that all scientific claims, design, data analysis, and raw text originate from the authors. Advanced automated textual processing software (specifically Anthropic Claude, OpenAI ChatGPT) was utilized exclusively for light code and light text editing. The entire manuscript and code base was reviewed by the authors, who maintain sole authorship and responsibility for the final text.

